# Brain endothelial cells exposure to malaria parasites links Type I interferon signalling to antigen presentation, immunoproteasome activation, endothelium disruption and cell metabolic adaptation

**DOI:** 10.1101/2022.11.26.518027

**Authors:** Abdul Muktadir Shafi, Ákos Végvári, Roman A Zubarev, Carlos Penha-Gonçalves

## Abstract

Cerebral malaria (CM) lethality is attributable to brain edema induction but the cellular mechanisms involving brain microvascular endothelium in CM pathogenesis are unexplored. Activation of the STING-INFb-CXCL10 axis in brain endothelial cells (BECs) is a prominent component of the innate immune response in cerebral malaria (CM) development in mouse models. Using a T cell-reporter system, we show that Type I IFN signaling in BECs exposed to *Plasmodium berghei*-infected erythrocytes (*PbA*-IE), functionally enhances MHC Class-I antigen presentation through gamma-interferon independent immunoproteasome activation and impacted the proteome functionally related to vesicle trafficking, protein processing, and folding and antigen presentation. *In vitro* assays show that Type I IFN signaling and immunoproteasome activation are also involved in the dysfunction of the endothelial barrier through disturbing gene expression in the Wnt/*ß*-catenin signaling pathway. We demonstrate that IE exposure induces a substantial increase in BECs glucose uptake while glycolysis blockade abrogates INFb secretion impairing immunoproteasome activation, antigen presentation, and Wnt/*ß*-catenin signaling. Metabolome analysis show that energy demand and production are markedly increased in BECs exposed to IE as revealed by enriched content in glucose and amino acid catabolites. In accordance, glycolysis blockade *in vivo* delayed the clinical onset of CM in mice. Our results unveiled that Type I IFN signaling and subsequent immunoproteasome activation in BECs contribute to enhanced antigen presentation and the impairment of the endothelial barrier function. We also show that Type I IFN signaling and its downstream effects are licensed by dramatic increase in glucose uptake impacting on energy metabolism pathways. This work substantiates the hypothesis that Type I IFN-immunoproteasome induction response in BECs contributes to CM pathology and fatality (1) by increasing antigen presentation to cytotoxic CD8+ T cells and (2) by promoting endothelial barrier dysfunction, favoring brain vasogenic edema.

**GRAPHICAL ABSTRACT:** 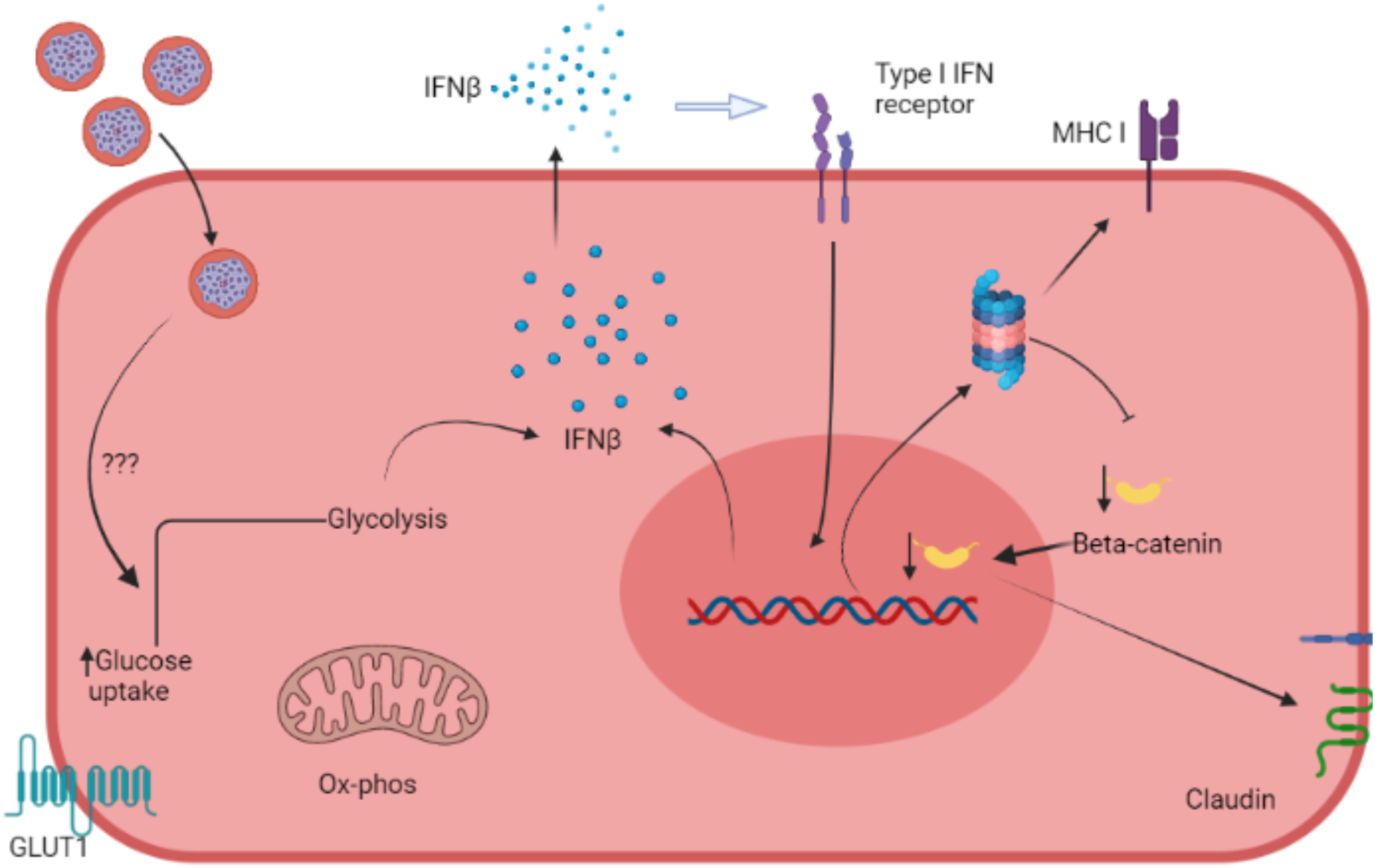

**AUTHOR SUMMARY:** Cerebral malaria is a severe clinical form of malaria infection that leads to respiratory arrest and death due to brain swelling. Disruption of blood vessels in the brain barrier is a hallmark of cerebral malaria development. However, the underlying pathological mechanisms are still unsettled. We explored the hypothesis that immune response of brain blood vessels to malaria infection is an initiator of inflammatory events leading to cerebral malaria. Our experiments unveiled that proinflammatory Type I interferon action increases the presentation of parasite molecules by brain blood vessel cells to cytotoxic immune cells and promotes brain vessel disruption. We found that these effects are determined by activation of protein degradation systems upon exposure of brain blood vessel cells to malaria parasite-infected erythrocytes. Our findings unveil a critical role of Type I interferon in brain blood vessels cells signaling in initiating immunopathology mechanisms associated to cerebral malaria development and suggest that blocking Type I interferon induction in the brain provides a path to prevent this disease.

## INTRODUCTION

Despite the deployment and rollout of state-of-the-art anti-malarial treatments, the malaria death toll reached 627000 cases in 2020 [1]. Malaria lethality is largely due to infection by *Plasmodium falciparum* in children and pregnant women developing severe forms of the disease [1]. Cerebral malaria (CM) is a neurological syndrome affecting mainly children. The accumulation of *Plasmodium falciparum* infected erythrocytes (IE) in the brain vasculature results in severe disease with patient mortality ranging from 15%-20% when effective anti-malarial treatment is available, and nearly 100% when patients are left untreated [2]. Adherence of IE to the brain endothelial cells triggers brain inflammatory responses and disruption of the blood-brain-barrier (BBB) leading to brain swelling, a hallmark of CM pathology [3]. However, the molecular mechanisms operating in brain endothelial cells to elicit brain immunopathology and BBB dysfunction are still unresolved.

Studies in *Plasmodium berghei* ANKA (*PbA*) mouse model of experimental CM (ECM) have identified a role for cytotoxic functions in CM-associated immunopathology. CD8 + T cell depletion and perforin or granzyme genetic ablation have shown that CD8^+^ T cells are critical in ECM pathogenesis. In 2020, Susan Pierce and colleagues [4] provided evidence of CD3^+^ CD8^+^ T cells in the luminal and perivascular spaces of brain blood vessels in fatal pediatric cerebral malaria cases from Malawi, mirroring observations in the ECM model [5]. Antigen presentation by class I molecules on brain endothelium dynamically regulates T-cell-mediated neuropathology in ECM [6]. Cytotoxic granzyme B, which is released when activated CD8^+^ T cells engage the cognate antigen-MHC class I complex, was detected inside and outside of CD8^+^ T cells lying on brain endothelial cells [7].

Innate immune responses involving induction of Type I IFN have been identified in multiple stages of malaria infection and are determinants of clinical disease trajectories [8]. In particular, Type I IFN receptor (IFNAR1) genetic variants were associated with cerebral malaria in children [9–11], and Type I interferon receptor (IFNAR1) signaling could enable effector functions of activated CD8^+^ T cells during ECM development [10]. We have shown that in the mouse brain parenchyma, brain endothelial cells are the main producers of IFNβ, a response determining the recruitment of leucocytes to the brain and immunopathogenesis of ECM [12,13]. Furthermore, the brain endothelium Type I interferon response may affect the interaction of endothelial-CD8+T cells or directly impact BBB permeability through unknown but critical cellular mechanisms.

The involvement of brain endothelial cells in effector phases of CM development is underlined by their ability to acquire and cross-present malaria parasite antigens both in human CM and ECM [4,14]. During *Pb*A infection, epitope-specific CD8+ T cells are induced in the spleen and migrate to the brain before the onset of neurological symptomatology [15]. Such epitope-specific CD8+ T cells are elicited by different murine malaria parasite strains but only the immune response triggered by *Pb*A causes neuropathology [15]. Notably, *Pb*A infection causes brain microvessels to cross-present parasite antigens, while non-ECM-causing parasites do not [15]. The precise mechanisms that are triggered by interaction with *Pb*A-IE in endothelial cells and that enable the cross-presentation of *Pb*A antigens are unexplored.

In the fatal cases of cerebral malaria, brain pathology shows either vasogenic edema or cytotoxic edema [16]. The tightness of BBB results from intercellular adherence provided by specialized tight junctions and by adherence junctions in the endothelia, which are regulated by *ß*-catenin-mediated classical Wnt signaling [17]. Alteration of the Wnt signaling leads to the downregulation of the tight junction proteins and increased BBB permeability [17–19]. It has been described that tight junction proteins and nuclear *ß*-catenin are decreased in hemorrhagic stroke-mediated BBB disruption [17–20]. However, it remains unclear whether IE-endothelial cell interactions may target Wnt signaling and promote BBB dysfunction during ECM development.

For several decades the metabolic relationships between parasites and infected erythrocytes were the focus of intense research leading to a better understanding of parasite biology, identification of anti-malarial drugs, and drug resistance mechanisms [21]. Recently, the hypothesis that the host’s systemic and cellular metabolism plays a role in the response to infection is shedding light on the disease mechanisms [22,23]. In *Plasmodium chabaudi* AJ murine model, perturbation of glycolysis using 2-deoxy-D-glucose (2-DG) reduced mice survival while administration of glucose had protective effects [24]. On the other hand, using *Pb*A infection, Wang et al (2018) showed that glycolysis modulation by competitive glucose analog 2DG protects mice from cerebral malaria. However, the role of endothelial cell metabolism in malaria has not been addressed. Endothelial cells are highly glycolytic, and glucose is a major contributor to cell energy demands. This is in line with the high expression of the glucose receptor GLUT1 found in many diseases affecting the central nervous system [18]. Stimulation of glycolysis promotes inflammation and shuttling of glucose towards the OXPHOS and the pentose phosphate pathway dampens inflammation [25].

Here, we investigated the downstream effects of Type I interferon signaling in brain endothelial cells (BECs) exposed to *Plasmodium berghei* ANKA infected erythrocytes (*Pb*A-IE) and their interplay with cellular metabolism. We found that the interaction of *Pb*A-IE with BECs elicits a Type I interferon response determining enhanced antigen processing, immunoproteasome activation, impaired Wnt*/ß*-catenin pathway signaling, and endothelial barrier disruption. This critical response of BECs is licensed by increased glucose consumption which impacts cellular energy metabolism pathways and is implicated in the initiation of ECM pathogenesis.

## RESULTS

### Type I interferon signaling controls endothelial antigen presentation and immunoproteasome induction

Interactions of brain endothelial cells (BECs) with IE are key in CM pathogenesis. To dissect molecular events resulting from these interactions we used an *in vitro* system where endothelial cell cultures derived from mouse brain vessels are exposed to *PbA-infected* erythrocytes (*Pb*A-IE). Malaria antigen cross-presentation was studied following a described method that uses a reporter T cell line recognizing the SQLLNAKYL *Pb*A peptide (derived from Glideosome-associated protein 50) in the context of mouse MHC-b haplotype (H-2D^b^) (Fig.1A) [26]. We quantified cross-presentation events to reporter T cells in primary BEC cultures. We found that BECs from IFNAR1 and IFNb KO mice show reduced efficiency in presenting malaria antigens (Fig. 1B). Medium supplementation with IFNβ rescued antigen presentation efficiency of IFNB KO BECs (Fig. 1C), indicates that the efficiency of malaria antigen presentation is determined by intracellular signaling through IFNAR1.

**Figure 1.**
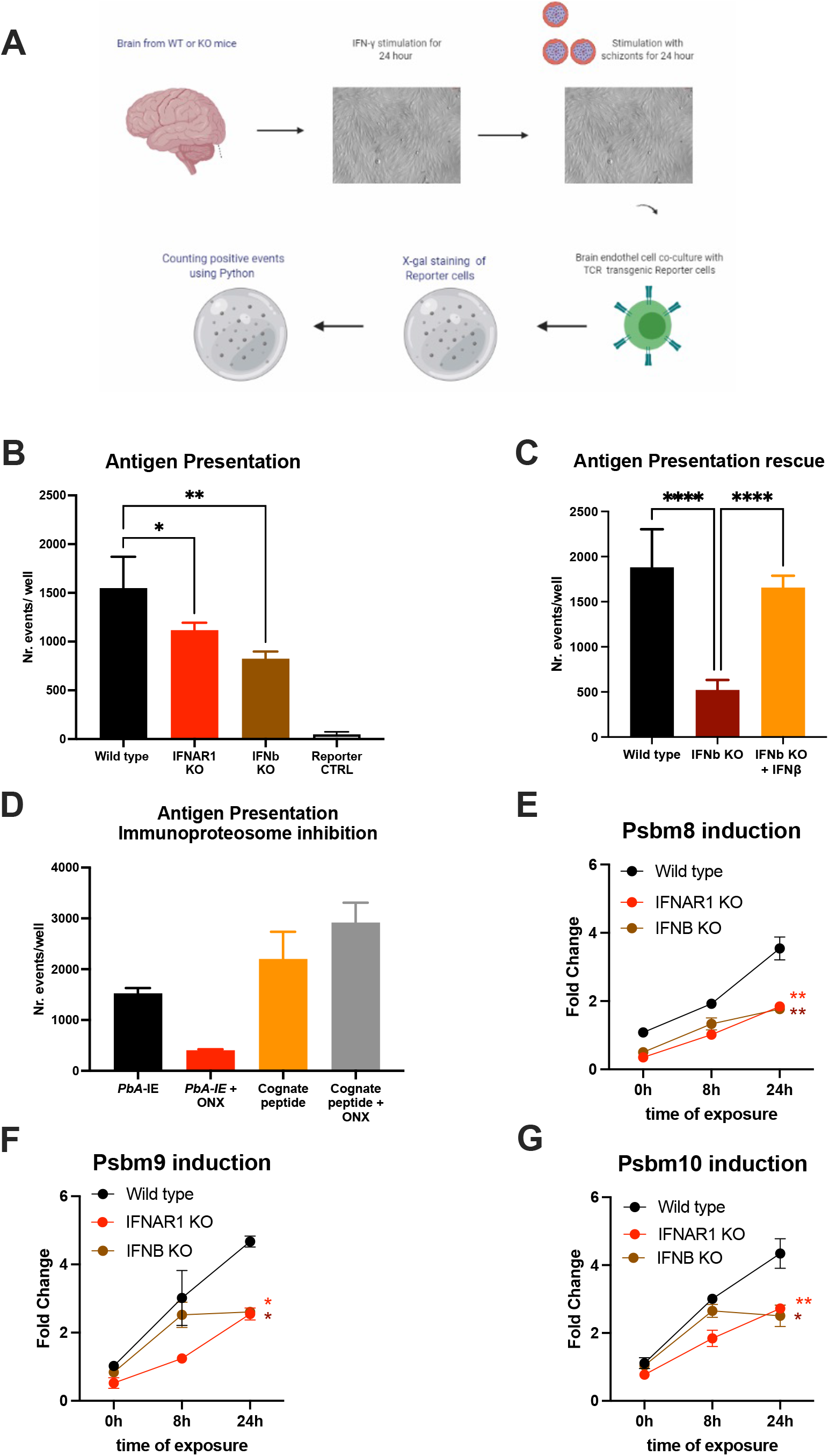
IFNAR1 signaling is a determinant of immunoproteasome-dependent antigen presentation in brain endothelial cells exposed to PbA-IE. **(A)** Diagram of in vitro antigen presentation assay depicting primary BEC cultured in presence of interferon-gamma (24h), exposed to synchronized Pba-infected erythrocytes (24h) and incubated with PbA antigen-specific reporter T-cell line subsequently stained to count antigen presentation events. **(B)** Counting antigen presentation events in BEC cultures from wild-type IFNAR1 and IFNB KO mice. **(C)** Cumulative results of two independent experiments counting antigen presentation events in BEC cultures from wild-type and IFNB KO mice in the presence or absence of IFNβ supplementation (80-120 μg/ml). **(D)** Cumulative results of two independent experiments counting antigen presentation events after exposing wild-type BEC cultures to PbA-IE or to reporter cell cognate peptide 10 μM with no supplementation with IFN-γ and in the presence or absence of immunoproteasome inhibitor ONX-0914 (300 nM)(ONX). Relative quantification of Psbm8 (E), *Psbm9* (F), and *Psbm10* (G) gene expression induction by qPCR in BECS of wild-type, IFNAR1 KO and IFNB KO mice exposed to PbA-IE in absence of interferon-gamma (fold change represents 2^ΔΔCT^ relative to wild-type at 0h). Statistics: P-value of ANOVA tests of antigen presentation results (black stars) or ANOVA tests of AUC in gene expression induction in IFNAR1 vs wild-type (red stars) and IFNB versus wild-type (brown stars). (*; p<0.05, **; p<0.01, ****; p <0.0001).

In response to inflammation or infection, the constitutive proteasome is replaced by an immunoproteasome, which is more efficient in processing endocytosed antigens for subsequent MHC I-restricted cross-presentation [27]. We examined *Plasmodium* antigen presentation by BECs under inhibition of immunoproteasome by ONX-0914 (PR-957) (ONX), a selective inhibitor of *Psmb8* [28]. We found that incubation with ONX 300 nM did not affect antigen presentation of MHC-I loaded cognate peptide but sharply decreased antigen presentation upon exposure to *PbA*-IE (Fig. 1 D). It is well-known that immunoproteasome subunits synthesis is induced by interferon-gamma (IFN-γ) and we asked whether immunoproteasome induction in BECs was controlled by Type I IFN signaling, in the absence of IFN-γ supplementation. We found that gene expression of three immunoproteasome subunits (*Psbm8, Psbm9*, and *Psbm10*) is induced in the course of exposure to *Pb*A-IE in wild-type BECs but was significantly lower in IFNAR1 KO and IFNb KO BECs (Fig. 1 E-F). Together these findings unveil that IFNAR1 signaling in BECs exposed to *Pb*A-IE operates the induction of immunoproteasome subunits synthesis that in turn contributes to the efficiency of intracellular *Plasmodium* antigens cross-presentation.

### Type I interferon signaling impacts protein processing and surface MHC class I molecules availability

Given that Type I IFN signaling was inducing the antigen processing machinery we asked whether it was also affecting the endothelial cell proteome. We performed a comparative proteome analysis in triplicate-independent samples of wild-type and IFNAR1 KO BECs exposed to *Pb*A-IE for 24 h, in the absence of IFN-γ supplementation. Notably, high-dimensional analyses (principal component analysis and orthogonal partial least squares discriminant analysis) of all mouse proteome features identified two clusters clearly distinguished according to the IFNAR1 genotype (Fig. 2A and B). Further, 207 protein features were identified as differentially expressed in wild-type and IFNAR1 KO BECs (univariate p value<0.05; fold change >1.2) (Fig. 2C). A more stringent selection of differentially expressed protein features (P<0.01) identified 34 proteins that were functionally grouped (Table S1). These analyses show that Type I IFN signaling significantly impacts endothelial proteome upon exposure to *Pb*A-IE and notably the expression of proteins related to antigen presentation, protein processing, and vesicle trafficking. Furthermore, flow cytometry analysis showed that IFNb KO BECs are not able to increase expression of surface expression of MHC class I molecules upon exposure to *Pb*A-IE (Fig. 2D). These results indicate that autocrine/paracrine Type I IFN secretion and subsequent signaling through IFNAR1 enhances the availability of MHC class I molecules for antigen presentation in BECs exposed to IE. Together our data uncover that Type I IFN signaling enhances BECs’ efficiency in presenting intracellular parasite protein antigens to CD8^+^T cells by acting on immunoproteasome activation, protein processing, vesicle trafficking and MHC clas,s I molecules availability.

**Figure 2.**
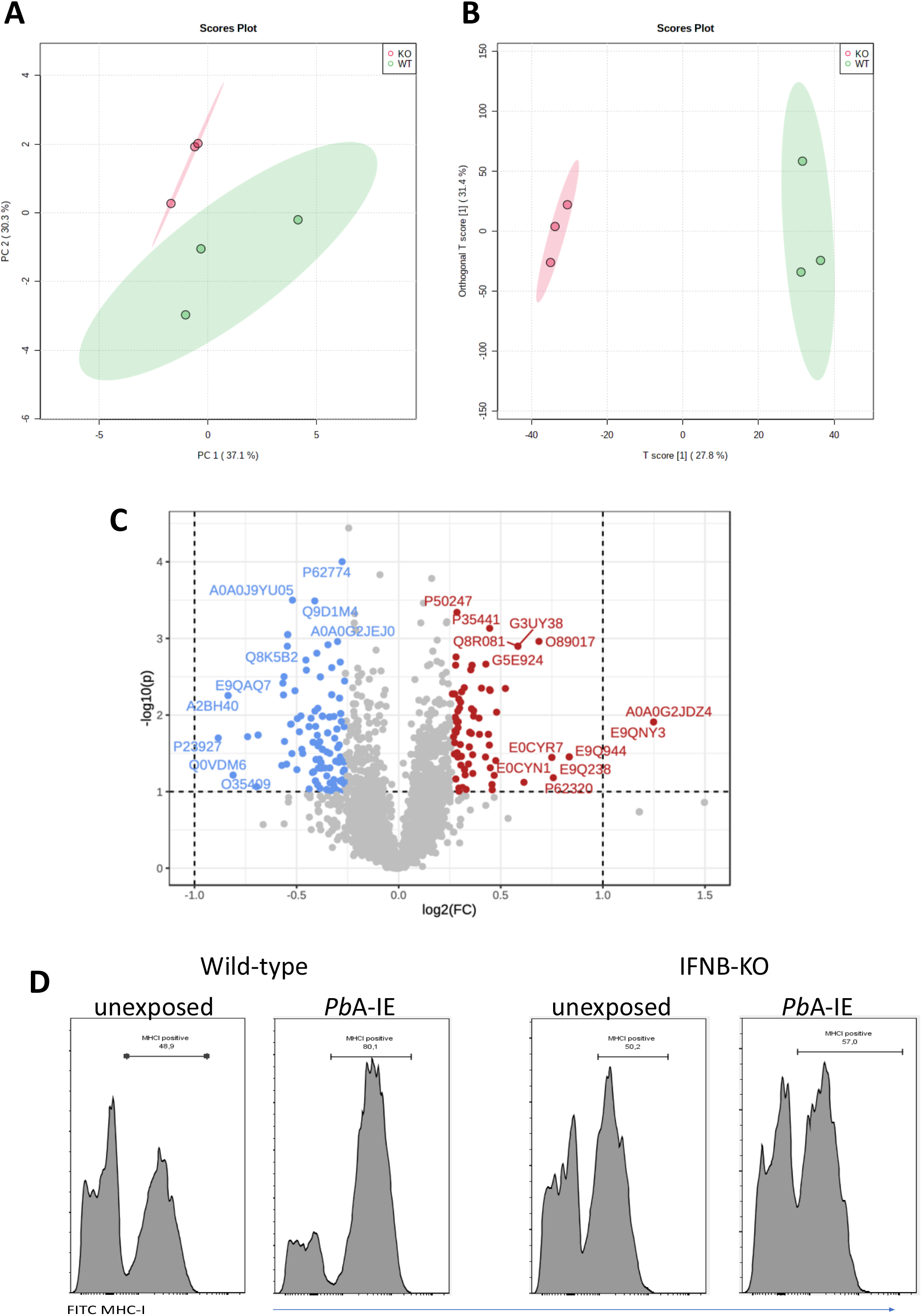
IFNAR1 signaling impact the proteome and MHC class I expression in BECs exposed to *Pb*A-IE. Proteome discrimination analysis of BECs cultures from IFNAR1KO and wild-type mice exposed to *Pb*A-IE. Principal component analysis score plot (PC1versus PC2) **(A)** and orthogonal partial least squares discriminant analysis (OPLS-DA) score plot **(B)** of all detected protein features in BECs samples (wild-type in green, IFNAR1 KO in red); explained variance is shown in brackets in axes titles. **(C)** Volcano plot highlighting differentially expressed protein features and showing fold change in IFNAR KO relative to wild-type (FC) > 1.2 (log2(FC) > 0.18, red; log2(FC) < - 0.18, blue) and t-test significance level, P<0.05. Differentially expressed protein features are identified by the respective SwissProt database reference. Proteome analysis is based on three independent samples per genotype, each sample representing BECs obtained from a pool of five mouse brains. **(D)** Flow cytometry histogram plots of surface staining of MHC class I molecules in BEC Interferon-beta deficient (IFNB KO) and wild-type mice, cultured in the absence or presence of *Pb*A-IE 24h before analysis. Results are representative of 2 independent experiments.

### IFNAR1 signaling and immunoproteasome contribute to endothelial barrier disruption

Type I IFN signaling in CD8+ T cells plays a critical role in cytotoxic functions that lead to BBB disruption during ECM development [10]. Exploring another angle, we examined whether Type I IFN signaling in BECs intrinsically contributed to the disruption of the brain endothelial barrier. BECs confluent cultures were challenged with *Pb*A-IE after trans-endothelial electric resistance (TEER), measured between the upper and lower chambers of a transwell system, increased and reached a high plateau that corresponds to the establishment of an endothelial barrier [29]. In wild-type BECs, TEER falls 60% by 24h after exposure to *Pb*A-IE indicating disruption of the endothelial barrier. In contrast, TEER showed a reduced decline in IFNAR1 KO BECs, indicating that the absence of IFNAR1 signaling offers partial resistance to endothelial barrier disruption upon *Pb*A-IE exposure (Fig. 3A). Proteome analysis corroborated that IFNAR1 signaling is associated with downregulation of an expressive number of proteins related to the dynamics of cytoskeleton organization that influence cell shape and motility possibly contributing to endothelial barrier disruption (Table S1).

**Figure 3.**
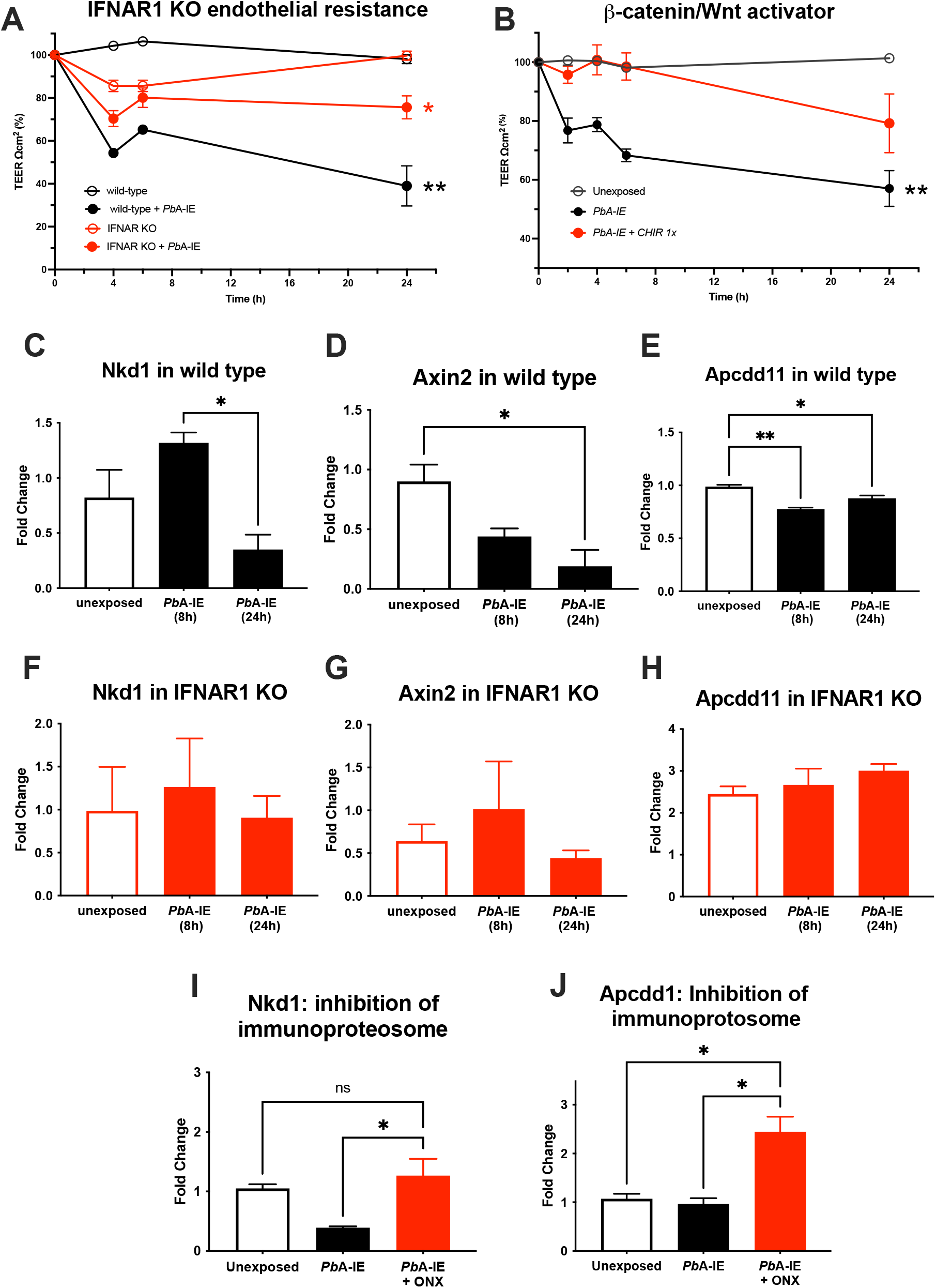
Wnt/*ß*-catenin signaling disturbances promoted by Type I interferon responses and immunoproteasome activation in response to *Pb*A-IE are associated with disruption of the brain endothelial barrier. Endothelial barrier integrity was monitored by electric resistance measurements of transwell BEC cultures at the indicated time points after exposure or not to *Pb*A-IE **(A and B)**. Kinetics of trans-endothelial electric resistance in wild-type or IFNAR1 KO BECs **(A)**, or in the presence or absence of (GSK) 3 inhibitor CHIR-99021(1 μm) wild-type BECs **(B)**. Gene expression of ***ß***-catenin target *genes Nkdd1, Axin2*, and *Apcdd1* was quantified by qPCR in wild-type **(C-E)** and IFNAR1 KO BECs **(F-H)** 24 h after exposure to *Pb*A-IE and presence or absence of immunoproteasome inhibitor ONX-0914 (ONX) (300 nM)**(I-J)**. Relative quantification of gene expression is represented as fold change (2^ΔΔCT^) using unexposed BECs as controls. Statistics: Significant results of ANOVA tests of AUC TEER in wild-type BECs exposed to PbA-IE versus unexposed (black stars) and in IFNAR1 KO BECs exposed versus unexposed (red stars). Significant results of pairwise comparisons in ANOVA tests in gene expression experiments are shown (*; p<0.05, **; p<0.01, ****; p <0.0001).

It is known that intracellular Wnt/*ß*-catenin pathway disturbances are associated with BBB dysfunction and are involved in the pathogenesis of several brain diseases [20]. We observed that *PbA-IE-induced* disruption of the endothelial barrier was prevented by BECs treatment with CHIR-99021, which effectively activates Wnt signaling by inhibiting glycogen synthase kinase (GSK) 3 which takes part of the *ß*-catenin destruction complex (Fig. 3B). Further, we found that expression of Wnt/*ß*-catenin signaling pathway genes (*Nkd1, Axin2*, and *Apcdd1*) was down-regulated in wild-type BECs upon exposure to *Pb*A-IE suggesting that Wnt/*ß*-catenin signaling was impaired (Fig. 3C-E). On the other hand, the expression of those target genes was not significantly disturbed in IFNAR1 KO BECs (Fig. 3 F-H). These results suggested that upon *Pb*A-IE challenge IFNAR1 signaling may contribute to endothelial barrier dysfunction by impairing Wnt/*ß*-catenin signaling.

Next, we explored the possibility that impaired transcription of Wnt/*ß*-catenin target genes resulted from increased immunoproteasome activation, which is known to promote the destruction of ubiquitinated *ß*-catenin [30]. We found that the immunoproteasome inhibitor (ONX) counteracts the downregulation of Wnt/*ß*-catenin signaling pathway genes induced by *Pb*A-IE (Fig. 3I-J). This suggests that immunoproteasome activation disturbs Wnt signaling possibly by acting on ubiquitinated *ß*-catenin. Cohesively, these findings suggest a mechanism of brain endothelium barrier disruption elicited by PbA-IE that involves Type I interferon-mediated immunoproteasome activity and consequent disturbance of WTN/B-catenin signaling.

Interestingly, gene expression of claudin 5, a tight junction protein that regulates paracellular ionic selectivity, was strongly down-regulated in wild-type BECs exposed to *Pb*A-IE (Fig. S1A). This effect was not restored in absence of IFNAR1 signaling neither was affected by immunoproteasome inhibition (Fig. S1B-C). Nevertheless, claudin 5 gene expression in the brains of infected mice was reduced as compared to non-infected animals (Fig. S1D). In alignment, proteins that are part of adherens-type cell-cell and cell-matrix junctions (e.g. thrombospondin-1, afadin, plectin, and phosphoglucomutase-like protein 5) are down-regulated by Type I IFN signaling in BECs exposed to *Pb*A-IE (Table S1). These results indicate that cell-cell junction disturbances evoked by *Pb*A-IE are in part dependent on Type I IFN signaling but suggest that additional mechanisms may play a role in endothelial barrier disruption caused by *Pb*A-IE.

### Glucose consumption promotes Type I interferon and associated responses in BECs exposed to *Pb*A-IE

Inflammatory stimuli have been shown to enhance glycolysis in endothelial cells [25]. We investigated whether glycolysis controls Type I IFN response in BECs exposed to *Pb*A-IE. To this end, we made use of a glucose analog 2-DG that competitively inhibits the production of glucose-6-phosphate from glucose, thereby blocking glycolysis. We found that adding 2-DG to BEC cultures abrogates IFN-ß secretion upon exposure to *Pb*A-IE (Fig. 4A), presumably impairing Type I IFN signaling-associated responses. We found that induction of immunoproteasome genes *Psmb8, Psmb9*, and *Psmb10* was decreased by 2-DG when BECs were exposed to *Pb*A-IE (Fig. 4B-D). Likewise, the efficiency of antigen presentation *in vitro* was affected when BEC cultures were incubated with 2-DG (Fig. 4E) which did not affect the ability to efficiently present cognate peptides loaded on surface MHC class I molecules (Fig. S2A). Notably, we performed *ex vivo* antigen presentation experiments using BECs from *Pb*A-IE infected mice and found that enhancement of antigen presentation by BECs exposed to *in vivo* infection was abrogated in 2-DG treated mice (Fig. 4F). In addition, we found that down-regulation of *Wnt/ß*-catenin target genes in *Pb*A-IE exposed BECs was reverted at some extent in presence of 2-DG (Fig. S2B). These results indicate that glucose metabolization is needed to license the observed Type I IFN signaling downstream effects, namely enhancement of antigen presentation and immunoproteasome activation.

**Figure 4.**
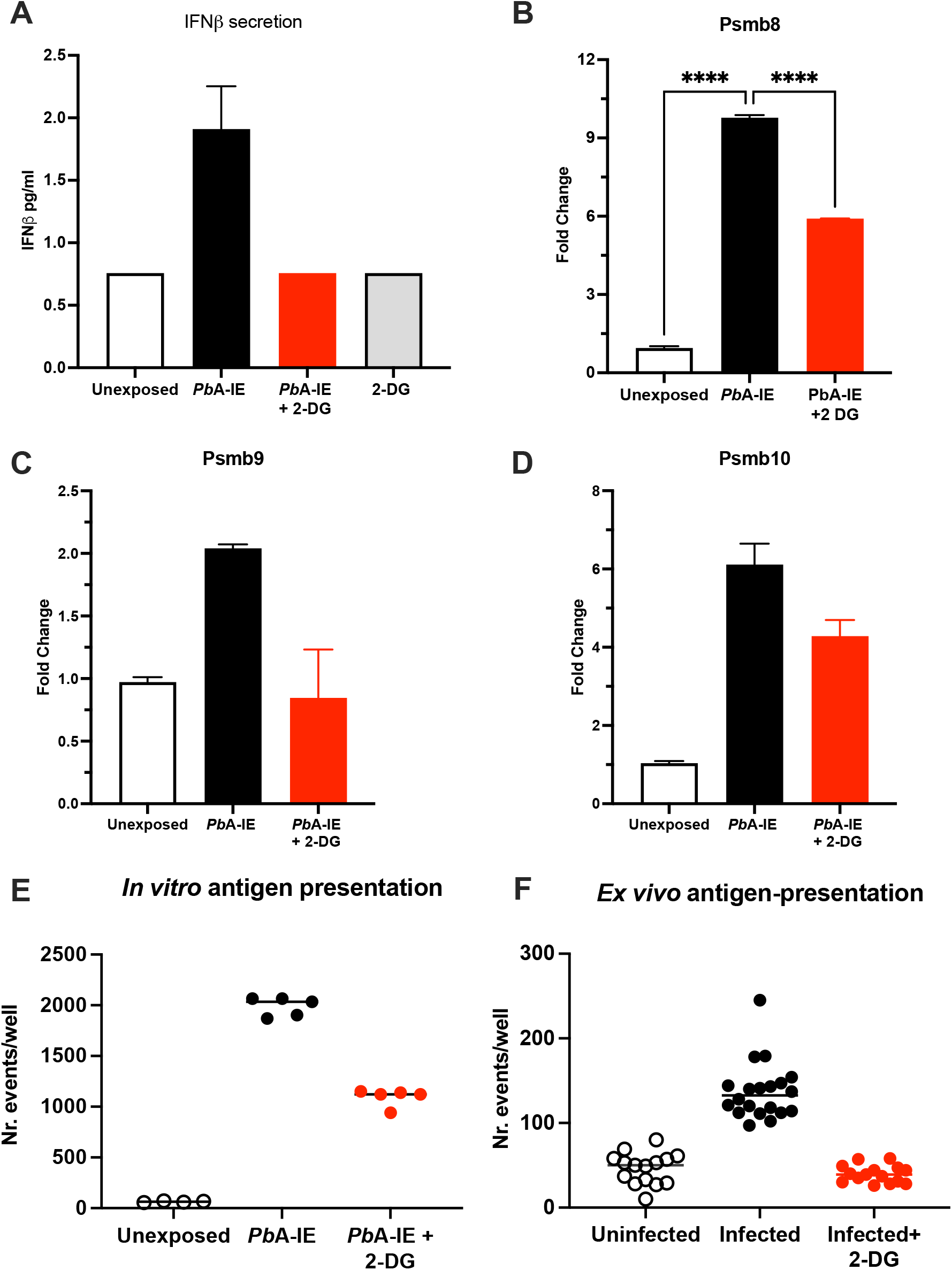
Glucose metabolism inhibition impairs IFN-ß secretion, immunoproteasome induction and antigen presentation in brain endothelial cells upon contact with *Pb*A-IE. Effects of inhibition of glycolysis by incubation of BEC cultures with 2-DG (10 mM) measured 24 h after exposure to *Pb*A-IE. **(A)** IFN-**ß** secretion was detected in culture supernatants by ELISA measurements BECs exposed or not to PbA-IE in the presence or absence of 2-DG (detection threshold 0.75 pg/mL). **(B-D)** Gene Expression of immunoproteasome genes Psmb8, Psmb9, and Psmb10 in BECs in the indicated conditions. Cumulative results of independent experiments are represented as fold change (2^ΔΔCT^) using unexposed BECs as controls. **(E)** Antigen presentation assays in BEC cultures unexposed and exposed to *Pb*A-IE *in vitro* in presence or absence of 2-DG, and in BECs from uninfected and infected mice treated or not with 2-DG **(F)**. Significant results of pairwise comparisons in ANOVA tests in gene expression experiments are shown (*; p<0.05, **; p<0.01, ****; p <0.0001).

### Metabolism perturbations in BECs exposed to *Pb*A-IE and in infected mice

To ascertain the effects of BEC exposure to *Pb*A-IE on glucose uptake we made use of 2-DG glycolysis inhibition and measured the amount of glucose in the culture supernatant as an indirect readout of glucose consumption by BECs. We found that 24h of BECs exposure to *Pb*A-IE increased glucose uptake signaled by a marked decrease of glucose in the cell culture medium which is significantly blocked in presence of 2-DG (Fig. 5A and Fig. S3A). Type I IFN signaling responses did not affect glucose uptake as BECs from IFNAR1 and IFNb KO mice exposed to *Pb*A-IE also showed marked glucose depletion from the cell culture medium (Fig. 5B). Nevertheless, proteome analysis in BECs exposed to PbA-IE shown that in presence of IFNAR1 signaling ATP enzymes related to ATP metabolism are up-regulated, namely ATP synthase subunit gamma and adenine phosphoribosyltransferase which critically contributes to AMP synthesis through the adenine salvage pathway (Table S2).

**Figure 5.**
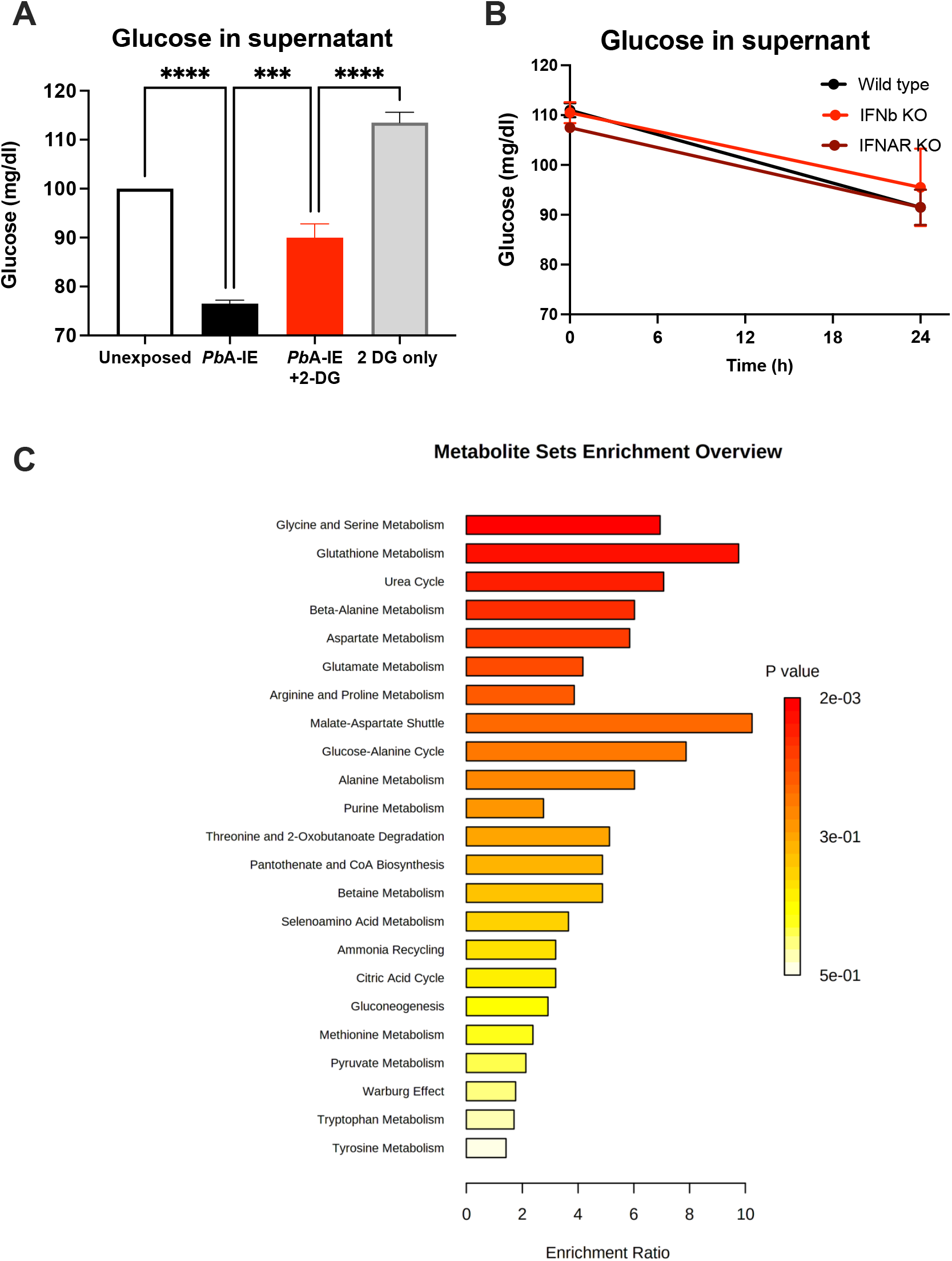
2-DG inhibits the uptake of glucose by brain endothelial cells exposed to *Pb*A-IE and increases the survival of infected mice. Measurement of glucose in supernatants of **(A)** BECs cultures after 24 h of exposure to *Pb*A-IE in the presence or absence of 2-DG (10 mM) and **(B)** in cultures from wild-type, IFNAR1 KO and IFNB KO BECs in presence of 2-DG. **(C)** Enrichment ratio (P-value) of metabolic pathways in wild-type BEC cultures exposed to *Pb*A-IE as compared to unexposed BEC cultures based on Multiple pathway targeted analysis data (MetaboAnalyst 5.0).

This led us to perform a multiple metabolic pathway targeted analysis comparing BECs exposed or not to *PbA-IE*, using HILIC - MS/MS as described in the methods section. We found that exposure to *PbA*-IE affected multiple pathways related to glucose metabolism which correlated with suggested increased energy demand and production (Fig. 5C) with a significant increase of individual metabolites such as pantothenate, creatine, and lactate (Table S2). We also noted found that the number of acylcarnitines was increased suggesting that the transport of fatty acids into mitochondria for energy production is enhanced following exposure to PbA-IE. In addition, we observed that amino acid metabolism was also affected in multiple pathways (Fig. 5C). Besides increased amounts of several amino acids (isoleucine, alanine, and aspartate) we also noted that intermediate metabolites related to amino acid catabolism (glutarate and 4-acetamidobutanoate) were increased. Unexpectedly, methylated amino acid derivatives such as trimethyllysine, dimethylarginine, and betaine (Table S2) were increased. Together with the observation that adenosylhomocysteinase is reduced in wild-type BECs exposed to PbA-IE (Table S1), these results suggest that the activated methyl cycle is affected by increased amino acid catabolism. Altogether these findings provide evidence that exposure to *Pb*A-IE strongly increased BECs glucose demand and impacted metabolic pathways and suggest that cellular downstream effects associated with Type I IFN signaling are dependent on increased glucose consumption.

### Glucose metabolism blockade in infected mice

Glucose metabolism mediates disease tolerance and resistance in cerebral malaria [22]. We tested whether the development of CM in the *PbA* infection mouse model was affected by the inhibition of cellular glucose uptake. We treated infected mice with one 2-DG injection (800 mg/kg) on day 4 when mouse BECs start expressing IFNβ and clinical symptoms of cerebral malaria are still incipient [13]. Daily clinical scoring of cerebral malaria revealed that 2-DG treatment delayed the clinical onset of neurological symptoms (Fig. 6A) and increased the survival time of infected mice compared to non-treated mice (Fig. 6B). As observed by others [31], 2-DG treatment immediately increased glucose blood levels indicating that cellular glucose uptake was reduced but, notably it did not reduce the amount of *Pb*A-IE as ascertained by the parasitemia level (Fig. S3B). This suggests that 2-DG treatment during infection exerted a protective effect on the brain tissue. In sum, these results indicate that increased glucose uptake and glycolysis elicited by *Pb*A-IE exposure affects Type I IFN mediated responses in BECs suggesting that beneficial effects of glucose blockade in the initiation of clinical CM may be linked to mechanisms operated by Type I IFN in BECs early in infection [13].

**Figure 6.**
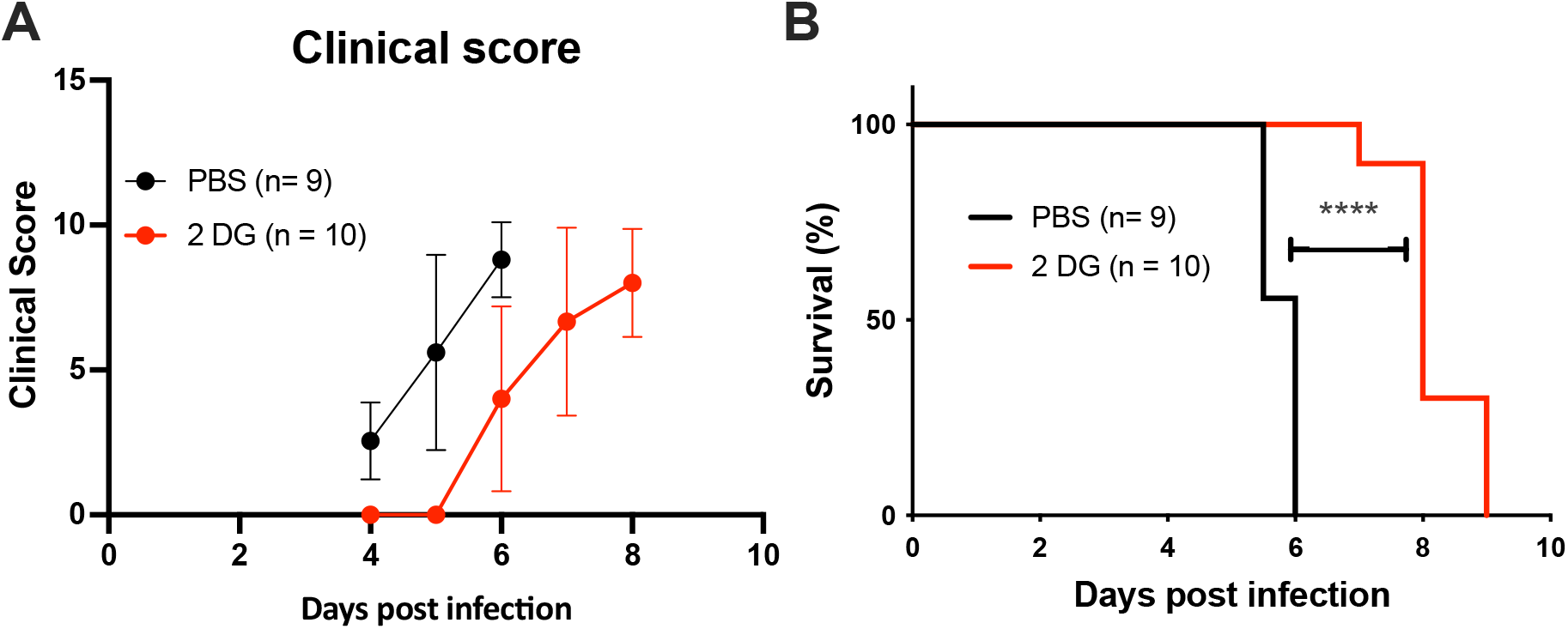
2-DG treatment increases the survival of infected mice. Clinical scoring **(A)** and survival curves **(B)** of mice infected with *Pb*A-IE and treated (2-DG) or not (PBS) with 2-DG (800 mg/kg) at day 4 post-infection. Significant results of pairwise comparisons in ANOVA tests **(A)**, and Log-rank (Mantel-Cox) test **(D)** are shown (**; p<0.01, ****; p <0.0001).

## DISCUSSION

This work links Type I IFN signaling in BECs exposed to *PbA-*IE to downstream cellular effects that eventually result in BBB damage. We found that Type I IFN increased the expression of immunoproteasome components and MHC class I molecules and, in interrelated association enhanced antigen presentation of parasite antigens to CD8+ T cells. In the course of the disease, this effect likely contributes to the cytotoxic effector functions of activated CD8+ T cells that target and damage the brain microvascular endothelium. In addition, immunoproteasome activation upon exposure to *PbA*-IE, conditioned disturbances of Wnt signaling that destabilized the endothelial barrier. This likely operates through imposing dysfunction on intercellular tight junctions, the first step towards increased BBB permeability which typically occurs in CM.

Remarkably, we found that increased glucose consumption in *PbA-*IE exposed BECs precedes Type I interferon response and its downstream effects. This uncovered that BEC’s innate immunity activation by *PbA-*IE is entangled with cellular metabolic alterations. Accordingly, we found that clinical ECM onset was delayed by blockade of glucose consumption, suggesting that increased glucose consumption is a precipitating factor in CM pathogenesis, possibly by licensing innate immunity activation of brain endothelial cells.

The pathogenic role of endothelium activation in the context of the innate immune response during malaria infection is catching increasing attention [12]. A large body of evidence links brain endothelium activation to the enhancement of adhesive properties underlying IE sequestration in the brain. ICAM-1, CD36, and EPCR have been identified as adhesion molecules that are up-regulated in activated BECs and act as counterparts of parasite molecules expressed at the IE surface [32]. The expression of endothelial adhesion molecules is induced by TNF during malaria infection, suggesting that the inflammatory milieu promotes changes in endothelium adhesivity [33,34]. IE sequestration has been considered a key event in CM pathology promoting local vascular inflammation and brain microcirculatory alterations [35]. Nevertheless, it is also clear that brain immunopathology events that lead to BBB disruption and brain edema play a determinant role in many CM cases [36].

Here, we investigated the effects of Type I IFN signaling that promote innate immunity functions in BECs and contribute to CM pathogenesis. Our approach unveiled the activation of cellular immunity mechanisms that promote antigen cross-presentation and are dependent on the autocrine action of IFN-ß. We have previously shown that IFNβ production by brain endothelial cells is a determinant of ECM development [13] and that CD8+ T cells required IFNAR1 expression to exert pathological effects in the brain and provoke ECM [10]. On the other hand, it has been demonstrated that BECs are induced to cross-present cytosolic malaria antigens to CD8+ T cells *in vitro* and *ex vivo* [15,26,37] Altogether, those results convey the notion that BEC’s Type I IFN response to IE autonomously provides two signals needed to engage activated cytotoxic CD8+ T cells namely, cross-presentation of parasite antigens in the context of MHC class I molecules, and IFNβ secretion. In this scenario, endothelial cells take a pivotal role in initiating adaptive immune responses that damage the BBB and lead to life-threatening cytotoxic edema.

On other hand, this autonomous Type I IFN signaling is also involved in weakening intercellular junctions of endothelial cells. This effect was associated with disturbances in the Wnt/*ß*-catenin signaling system. It has been reported that in human brain endothelial cells, *ß*-catenin activation upon exposure to *Plasmodium falciparum*-IE was interrelated with angiotensin II signaling pathways [38]. We found that Type I IFN signaling-dependent disturbances in the Wnt/*ß*-catenin system, were associated with endothelial junctions dysfunction. This provides initial evidence of an autocrine/paracrine mechanism leading to dysfunction of the endothelial barrier acting in absence of other cellular or molecular components of the immune system. Such a mechanism would allow extravasation of intravascular fluid to the perivascular space, a critical step for vasogenic brain edema that is a frequent cause of death in subjects with cerebral malaria [37,39].

It is well known that immunoproteasome induction acts in the processing of cytosolic antigens in preparation for their presentation in the context of MHC I molecules [40]. We found that immunoproteasome gene expression was induced in BECs exposed to *PbA-*IE. Such immunoproteasome activation likely contributes to the observed enhancement of antigen presentation efficiency in corroboration with previous findings that *Plasmodium* antigens are processed via a cytosolic pathway rather than the vacuolar pathway [15,41]. The *ß*-catenin cytoplasmic pool is tightly regulated via the ubiquitin-proteasomal pathway [19,42] and we found that gene expression driven by Wnt/*ß*-catenin in BECs exposed to *PbA*-IE is partially recovered when the immunoproteasome is inhibited. Likewise, proteome analysis shows that the expression of several proteins related to cytoskeleton organization and cell-cell junctions was altered by Type I IFN signaling. This suggests that immunoproteasome activation and other cellular processes induced by Type I IFN signaling indirectly affect the stability of endothelial intercellular junctions which partially operate under the control of Wnt/*ß*-catenin signaling.

Moreover, we show that immunoproteasome induction in BECs was dependent on Type I IFN signaling but in contrast to established mechanisms did not require exogenous IFN-γ [43]. This unexpected finding is only paralleled by observations that during hepatitis C virus infection immunoproteasome induction in the liver occurs in a Type I IFN-dependent manner [44]. The precise mechanisms of immunoproteasome induction by Type I IFN are still unclear but further research may shed light on how Type I IFN may operate in other non-immune cells to engage adaptive immune responses.

We found that IFNβ secretion in *PbA-IE* exposed BECs was preceded by and dependent on increased glucose consumption. Likewise, the blockade of glycolysis affected the downstream effects of Type I IFN signaling including, immunoproteasome induction, enhancement of antigen presentation, and down-regulation of Wnt signaling gene expression. It is well-known that in response to inflammation, cells shift glucose metabolism toward glycolysis [45–47].

Remarkably, the sharp increase in glucose uptake in BECs upon contact with IE did not depend on host extra-endothelial inflammatory or metabolic stimuli. It is interesting that besides impacting energy metabolism IE exposure also increased amino acid catabolism and methylation. It is worth exploring whether these perturbations in amino acid metabolism are related to the activation of immunoproteasome and enhanced antigen presentation pathway. Furthermore, amino acid metabolism may play a role in controlling the redox state cells exposed to *Pb*A-IE. The synthesis and catabolism of amino acids are interwoven into the redox homeostasis of the cell. The malate–aspartate shuttle, as well as moving NADH between the cytosol and the mitochondrial matrix, also moves the amino acids glutamate and aspartate between the two compartments and is functionally connected to the TCA cycle.

Our results also indicate that increased glucose uptake elicited by *Pb*A-IE infection is critical to the induction of IFNAR1 signaling effects in BECs that in turn precipitate pathogenesis mechanisms operating in the initiation of clinical CM in the mouse [13] The IE components and IE-BECs interaction events that lead to increased glucose uptake by BECS remain to be identified.

Overall, this work provides evidence that Type I IFN response in BECs exposed to IE is preceded by glucose metabolism alterations and is a driver of multiple effector mechanisms that likely underlie the two main pathogenesis determinants of CM fatality: brain immunopathology and vasogenic edema.

## MATERIALS AND METHODS

### Animals

All animal procedures were conducted according to national (Portaria 1005/92) and European regulations (European Directive 86/609/CEE) and were approved by the Instituto Gulbenkian de Ciência Ethics Committee (Project number A05.2020) and the Direcção-Geral de Alimentação e Veterinária, the national authority for animal welfare. Mice were housed and bred in the facilities of Instituto Gulbenkian de Ciência. C57BL/6 mice (wild-type) and *Ifnar1*^*-/-*^ (IFNAR KO) mouse strains were obtained from the Instituto Gulbenkian de Ciência facilities. *Ifnβ1*^−/−^ (IFNβ KO) mouse strain was obtained from the TWINCORE, Centre for Experimental and Clinical Infection Research, Hanover, Germany.

### Murine cerebral malaria model

Wild-type or KO mice were infected with 10^6^ *Pb*A-GFP (intraperitoneal route) obtained from blood-frozen stocks of infected C57BL/6 mice. Parasitemia in infected mice was determined by flow cytometry analysis as GFP+ erythrocytes (LSRFortessa™ X-20 cell analyser, BD Biosciences, and FACSDiVa software version).

Cerebral Malaria progression was evaluated with a 10-point clinical scoring method covering behavioral, visual, gait, and motor functions. Animals were scored on five attributes, observational and task-based, interactions/reflex, cage grasp, visual placing, gait/posture/appearance, and capacity to hold their body weight on a baton. Every attribute was scored from 0 to 2, where 0 represents normal and 2 represents strong impairment [48]

### Brain endothelial cell (BEC) cultures

Mouse brains were harvested in an ice-cold HBSS medium (Biowest). Brains were minced using a scalpel blade and homogenized by passing through a 23G needle. Homogenates were mixed with an equal volume of 30% dextran (Sigma) and centrifuged at 10000g for 15 minutes. This led to the isolation of brain microvessel pellets, which were suspended in DMEM and passed through a 40u strainer. Brain micro-vessels were retrieved by backflushing with 5 ml of MD131 and were incubated for 2 h at 36.5 ºC in an orbital shaker with a digestion cocktail containing 1mg/ml of collagenase type IV (Millipore), 10 μg/ml DNAse (Roche) and 2% FBS. Digested brain microvessels were suspended and cultured in 6, 24 and 48 well-plates in EGM™-2 Endothelial Cell Growth Medium-2 BulletKit™ (LONZA) supplemented with GlutaMax (ThermoFisher) and 10% of FBS, and containing 4 μg/ml puromycin. After 4 days, puromycin containing medium were changed and every 4 days cells were fed EGM™-2 Endothelial Cell Growth Medium-2 BulletKit™ (LONZA) supplemented with GlutaMax (ThermoFisher) and 10% of FBS. When cultures reached 50-80% confluence (∼14 days, 95% CD31+ by flow cytometry analysis), the cells were used for experiments.

### Preparation of synchronized infected erythrocytes (*Pb*A-IE)

Blood was collected by cardiac puncture with a heparinized needle from *Pb*A infected mice when parasitemia reached 10% and harvested in 5 ml of RPMI1640 culture medium. Blood was centrifuged at 450g for 8 min and the supernatant was discarded. Erythrocytes were resuspended in RPMI1640 containing 20% FBS in a 250 ml cell culture flask and flushed with a gas-mixture of 5% CO2, 5% O2 and 90% N2. The sealed flasks were incubated in an orbital shaker at 3forring 16 h to synchronize infected erythrocytes in the schizont stage (*Pb*A-IE).

### Cross presentation assays

*In vitro* cross-presentation assays were performed following a previously described protocol [26] briefly described below., Confluent BEC cultures were stimulated with IFNγ (10 ηg/ml), unless stated otherwise. Twenty-four hours after stimulation, cells were stimulated with 2-3 × 10^6^ *Pb*A-IE. After another 24 hours, the medium was withdrawn and the BEC culture was washed with RPMI complete medium and incubated with 6 × 10^4^ reporter cells carrying a TCR transgene that specifically recognizes the PbA peptide (SQLLNAKYL). After overnight incubation, reporter cells were resuspended and added to 96 well filter plate and stained with X-gal. For specific experiments, the medium was supplemented with immunoproteasome inhibitor ONX-0914 (300 nM) or with the glucose analog, 2-DG (10 mM), added 24 hours before cell harvesting.

For *ex-vivo* cross-presentation, brain micro-vessels were isolated from infected and uninfected mice at 6 days after infection (when the wild-type mice show signs of ECM). After isolation, the micro-vessels were passed through a 40 μm strainer and backflushed with 2 ml PBS before incubation with the digestion cocktail (1mg/ml of collagenase type IV (Millipore), 10 μg/ml DNAse (Roche) and 2% FBS) for 1.5 hours. After digestion, microvessels from each brain were distributed per five 96 well filter plates and 3x10^4^ reporter cells were added to each well. 16-18 hours after co-incubation, reporter cells were stained for X-gal.

After staining, the images were captured using a customized stereoscope, assembled with a TCZRS series, Bi-telecentric zoom lenses with motorized control (https://www.opto-e.com/products/8x-bi-telecentric-zoom-lenses-with-motorized-control#Matrix), and RT-mvBF3-2051a, USB3 Vision camera with Sony Pregius CMOS sensor IMX264. Images were captured at 1x magnification with circular illumination. For *ex-vivo* assays, images were processed in ImageJ, and positive events were counted using the find maxima routine. For *in vitro* image processing, event counts were obtained by using an image analysis dedicated script that was implemented to selectively exclude signals from schizonts remnants (red) and only include reporter cells signals (blue).

### Gene expression

BECs were recovered from culture wells. For specific experiments, the medium was supplemented with immunoproteasome inhibitor ONX-0914 (300 nM) or with the glucose analog, 2-Deoxy-d-glucose (2-DG) (10 mM), added 24 h before cell recovery.

cDNA was prepared from each culture well, using a Cells-to-CT kit (Thermo Fisher Scientific). qRT-PCR reactions using standard protocols were performed in ABI QuantStudio-LDA with the following Taqman assays: Mm00440207_m1(Psmb8), Mm00479004_m1 (Psmb9), Mm00479052_g1 (Psmb10), Mm01257559_m1 (Apcdd1), Mm00443610_m1 (Axin2), Mm00727012_s1 (Cld5), Mm00438656_m1 (Edn1), Mm00471902_m1 (Nkd1) and Mm00607939_s1(ACTB), as an internal control. Relative gene expression was quantified using the 2^ΔΔCT^ method.

### Sample preparation for Proteomics

Brain endothelial cell cultures from five mouse brains of IFNAR1 KO and wild-type mice were prepared in six-well plates and exposed to 5 x10^6^ *Pb*A-IE/well, in the absence of IFN-γ supplementation. Twenty-four hours after stimulation, un-phagocytosed schizonts were washed out and BMECs were lysed using RIPA lysis buffer followed by precipitation with acetone. For precipitation, 4 x volume chilled acetone was added to the lysate and incubated at 20° overnight. After that, the sample was centrifuged at 15000g for 15 min and protein pellets were isolated and sent for mass spectrometric analysis. Cell pellets were solubilized with 10 μL of 8M urea and 10 μL of 1% Protease MAX (Promega) in 100 mM Tris-HCl, pH 8.5 following sonication in a water bath for 5 min. Following 79 μL of 50 mM Tris-HCl was added together with 1 μL of 100x protease inhibitor cocktail (Roche) before the probe sonicated with VibraCell probe (Sonics & Materials, Inc.) for 40 s, with pulse 2/2, at 20% amplitude. Protein concentration was measured by BCA assay (Thermo Fisher Scientific).

Sample processing and mass spectrometric analysis was performed at the Proteomics Biomedicum (Karolinska Institutet, Stockholm). Briefly, an aliquot of 15 μg sample was transferred to a new tube and equalized with 50 mM Tris-HCl to a total volume of 90 μL. Proteins were reduced with the addition of 3 μL of 250 mM dithiothreitol (Sigma) at 37°C for 45 min and alkylated with 4.5 μL of 500 mM chloroacetamide for 30 min at room temperature in the dark. Sequencing grade trypsin (Promega) was added in an enzyme-to-protein ratio of 1:50 (1.5 μL of 0.2 μg/μL) and digestion was carried out overnight at 37°C. The digestion was stopped with 5 μL cc. formic acid (FA), incubating the solutions at RT for 5 min.

The sample was cleaned on a C18 Hypersep plate with 40 μL bed volume (Thermo Fisher Scientific), and dried using a vacuum concentrator (Eppendorf). Biological samples were labeled with TMT-6plex reagents in random order adding 120 μg TMT-reagent in 30 μL dry acetonitrile (ACN) to each digested sample resolubilized in 70 μL of 50 mM triethylammonium bicarbonate (TEAB) and incubating at room temperature (RT) for 2 h. The labeling reaction was stopped by adding 11 μL of 5% hydroxylamine and incubating at RT for 15 min before combining them in one vial.

### Liquid Chromatography-Tandem Mass Spectrometry Proteome Data Acquisition

Peptides were reconstituted in solvent A (2% ACN/0.1% FA) and approximately, two μg samples (2/12 μL) were injected on a 50 cm long EASY-Spray C18 column (Thermo Fisher Scientific) connected to an Ultimate 3000 nanoUPLC system (Thermo Fisher Scientific) using a 90 min long gradient: 4-26% of solvent B (98% ACN/0.1% FA) in 90 min, 26-95% in 5 min, and 95% of solvent B for 5 min at a flow rate of 300 nL/min. Mass spectra were acquired on a Q Exactive HF hybrid quadrupole orbitrap mass spectrometer (Thermo Fisher Scientific) ranging from *m/z* 375 to 1700 at a resolution of R=120,000 (at *m/z* 200) targeting 5x10^6^ ions for a maximum injection time of 80 ms, followed by data-dependent higher-energy collisional dissociation (HCD) fragmentations of precursor ions with a charge state 2+ to 8+, using 45 s dynamic exclusion. The tandem mass spectra of the top 18 precursor ions were acquired with a resolution of R=60,000, targeting 2x10^5^ ions for a maximum injection time of 54 ms, setting quadrupole isolation width to 1.4 Th and normalized collision energy to 33%.

### Proteome Data Analysis

Acquired raw data files were analyzed using Proteome Discoverer v2.5 (Thermo Fisher Scientific) with Mascot Server v2.5.1 (Matrix Science Ltd., UK) search engine against mouse protein database (SwissProt). A maximum of two missed cleavage sites were allowed for full tryptic digestion, while setting the precursor and the fragment ion mass tolerance to 10 ppm and 0.02 Da, respectively. Carbamidomethylation of cysteine was specified as a fixed modification, while TMT6plex on lysine and N-termini, oxidation on methionine as well as deamidation of asparagine and glutamine were set as dynamic modifications. Initial search results were filtered with 5% FDR using the Percolator node in Proteome Discoverer. Quantification was based on the TMT-reporter ion intensities. Proteome discrimination of three samples per genotype (IFNAR1 KO or wild-type) was obtained using all proteome features with high-dimensional analysis methods; principal component analysis (PCA) and orthogonal partial least squares discriminant analysis (OPLS-DA). Volcano plots were used to identify proteome features differentially represented in IFNAR1 KO BEC samples.

### Multiple pathway targeted analysis

Cell pellets were pre-extracted and homogenized by the addition of 1000 μL of MeOH: H2O (4:1), in the Cryolys Precellys 24 sample Homogenizer (2 × 20 seconds at 10000 rpm, Bertin Technologies, Rockville, MD, US) with ceramic beads. The bead beater was air-cooled down at a flow rate of 110 L/min at 6 bar. Homogenized extracts were centrifuged for 15 minutes at 4000 g at 4°C (Hermle, Gosheim, Germany). The resulting supernatant was collected and evaporated to dryness in a vacuum concentrator (LabConco, Missouri, US). Dried sample extracts were resuspended in MeOH: H2O (4:1, v/v) according to the total protein content. Media samples (25 μL) were extracted with 100 μL of MeOH. Homogenized extracts were centrifuged for 15 minutes at 4000 g at 4°C (Hermle, Gosheim, Germany). The resulting supernatant was collected and injected into the LC-MS system. BCA Protein Assay Kit (Thermo Scientific, Massachusetts, US) was used to measure (A562nm) total protein concentration (Hidex, Turku, Finland).

Extracted samples were analyzed by Hydrophilic Interaction Liquid Chromatography coupled to tandem mass spectrometry (HILIC - MS/MS)1,2 in both positive and negative ionization modes using a 6495 triple quadrupole system (QqQ) interfaced with 1290 UHPLC system (Agilent Technologies). Pooled QC samples (representative of the entire sample set) were analyzed periodically (every 4 samples) throughout the overall analytical run in order to assess the quality of the data, correct the signal intensity drift and remove the peaks with poor reproducibility (CV > 30%) [49–51]

### Flow Cytometry

After the confluence of the BMEC, cells were detached using TrypLE™ Express Enzyme (1X), no phenol red, and stained with anti-CD31 (APC, Anti-Mouse) (BD Pharmingen™) MHCI (A488, Anti-Mouse) and propidium iodide as per the standard flow cytometry protocol and analyzed thereafter (LSRFortessa™ X-20 cell analyzer, BD Biosciences, and FACSDiVa software version).

### Measurements of transendothelial electrical resistance TEER

BECs were grown to confluence in VWR Tissue Culture Plate Inserts 24 well PET membrane, 8μm. Trans-endothelial electric resistance (TEER) was recorded using Millicell ERS-2 Voltohmmeter according to the manual. A detailed description of this methos has been published elsewhere [29]. Once a stable measurement of resistance was achieved, BECs in the inserts were exposed to 1.5 × 10^6^ freshly prepared *Pb*A-IE or left unexposed and treated or not with GSK 3 inhibitor CHIR-99021(1 μm) for 24 h. Electric resistance was monitored at the indicated time points.

### Glycolysis inhibition and glucose measurements

For *in vivo* glycolysis inhibition experiments mice were injected intra-peritoneally with 2-DG (800 mg/kg) at day 4 four days after infection with *Pb*A-IE. The timing of injection was chosen according to evidence that IFNb is expressed in the brain by day 4 after infection [13]. For measurement of glucose uptake, BECs were cultured in 48 well culture plates and exposed to 1.5 to 3 × 10^6^ schizonts for 24 h in presence of absence of 2-DG (10 mM). The supernatant was collected at specific time points during this period and centrifuged at 10000g for 2 min. Glucose was measured by @Accucheck glucose measurement kit. To ascertain whether glucose consumption was not attributable to *Pb*A-IE a set of experiments, where after 24-h exposure of the BECs to *Pb*A-IE, the supernatant containing IE was discarded and BECs properly washed to avoid any leftover of the IE. Thereafter fresh medium was dispensed in each well for 24 h and glucose was measured in the collected media (Fig. S3).

### Statistics

Calculations of ANOVA tests, area under the curve (AUC), and Log-rank (Mantel-Cox) test were performed using Prism 9 for macOS from GraphPad Software, LLC.

## ACKNOWLEDGMENTS

We wish to thank Dr. Laurent Renia, NTU, Singapore that kindly provided us the transgenic reporter T cell line we used in antigen presentation assays. We thank dr. Ulrich Kalinke, from TWINCORE, Centre for Experimental and Clinical Infection Research, Hannover, Germany for making IFNb ko mice available to us. We thank Abdul Muntakim Rafi for his help with image analysis and the technical assistance from the imaging facility at the IGC. This work was developed with the support from CONGENTO LISBOA-01-0145-FEDER-022170, co-financed by FCT (Portugal) and Lisboa2020, under the PORTUGAL2020 agreement (European Regional Development Fund). We thank Drs. Teresa Pais and Nádia Duarte for the critical reading of the manuscript.

## SUPPLEMENTARY INFORMATION

**Table S1:** Differential protein expression in BECs from IFNAR1 KO and wild-type mice upon *in vitro* exposure to *PbA*-IE. Functional grouping of selected differentially expressed protein features.

| Grouping | Protein | Feature ID | log<sub>2</sub>(FC) | -log<sub>10</sub>(p) | Obs |
| --- | --- | --- | --- | --- | --- |
| **Antigen presentation** |  |  |  |  |  |
|  | DC-STAMP domain-containing 2 | A0A140LIJ0 | -0.57 | 2.40 |  |
|  | H-2 class I histocompatibility antigen, L-D alpha chain | P01897 | -0.26 | 2.48 |  |
|  | Legumain | O89017 | 0.69 | 3.01 |  |
| **Protein trafficking and folding** |  |  |  |  |  |
|  | Multiple coagulation factor deficiency protein 2 homolog | Q8K5B2 | -0.55 | 2.93 |  |
|  | Prefoldin subunit 3 | Q3TIR6 | -0.33 | 2.32 | 2 features |
|  | Calumenin | Q6XLQ8 | -0.33 | 2.61 | 2 features |
|  | GTPase HRas | COH5X4 | 0.26 | 2.33 |  |
|  | Vacuolar protein sorting-associated protein 26A | A0A1W2P7Z9 | 0.37 | 2.08 |  |
| **Vesicle trafficking** |  |  |  |  |  |
|  | Protein S100-A10 | P08207 | -0.20 | 2.68 |  |
|  | Y-box-binding protein 1 | P62960 | -0.19 | 2.70 |  |
|  | Ras-related protein Rab-18 | P35293 | 0.13 | 2.62 |  |
|  | Protein transport protein Sec31A | Q3UPL0 | 0.19 | 2.87 |  |
|  | Annexin A6 | F8WIT2 | 0.21 | 2.52 | 2 features |
|  | Protein transport protein Sec31A | S4R2A9 | 0.23 | 3.12 |  |
|  | EH domain-containing protein 2 | Q8BH64 | 0.32 | 2.38 |  |
| **Cytoskeleton/cell shape and motility** |  |  |  |  |  |
|  | WD repeat-containing protein 1 | A0A0J9YU05 | -0.52 | 3.48 |  |
|  | Myosin regulatory light polypeptide 9 | Q9CQ19 | -0.50 | 2.35 |  |
|  | Actin, gamma-enteric smooth muscle | A0A0U1RQ96 | -0.45 | 2.74 |  |
|  | Transgelin | A0A1L1STN8 | -0.45 | 2.59 | 2 features |
|  | Myristoylated alanine-rich C-kinase substrate | P26645 | -0.40 | 2.79 |  |
|  | Vimentin | P20152 | -0.29 | 2.69 |  |
|  | Myosin light polypeptide 6 | Q60605 | -0.27 | 2.72 | 4 features |
|  | laminin subunit alpha-4 | P97927 | 0.29 | 2.20 |  |
|  | Laminin subunit beta-2 | Q61292 | 0.45 | 2.33 |  |
| **Extracellular matrix/cell adhesion** |  |  |  |  |  |
|  | Zyxin | A0A0N4SVD2 | -0.54 | 3.07 | 3 features |
|  | Galectin-1 | P16045 | -0.35 | 2.86 |  |
|  | PDZ and LIM domain protein 5 | A0A0G2JEJ0 | -0.30 | 3.00 |  |
|  | Phosphoglucomutase-like protein 5 | Q8BZF8 | 0.28 | 2.75 |  |
|  | Ras-interacting protein 1 | Q3U0S6 | 0.30 | 2.28 |  |
|  | Afadin | A0A668KLE6 | 0.36 | 2.09 |  |
|  | Heparan Sulfate Proteoglycan 2 | E9PZ16 | 0.36 | 2.80 | 3 features |
|  | Serine protease HTRA1 | Q9R118 | 0.41 | 2.31 |  |
|  | Thrombospondin-1 | P35441 | 0.45 | 3.13 |  |
| **Metabolism** |  |  |  |  |  |
|  | ATP synthase, H<sup>+</sup>-transporting, F1 complex |  | -0.30 | 2.01 | 7 features |
|  | Adenine phosphoribosyltransferase |  | -0.40 | 2.08 |  |
|  | Adenosylhomocysteinase |  | 0.29 | 3.34 |  |

FC represents KO/wild-type relative abundance. Shown are protein features with P value <0.01 (-log<sub>10</sub>(p)<2) and log<sub>2</sub>(FC) > 0.18, red; log<sub>2</sub>(FC) < - 0.18, blue. For proteins with more than one significant feature, the feature with largest FC effect is shown.
Proteins identified from respective feature ID in the SwissProt database. FC represents KO/wild-type rate. Shown are protein features with P value <0.01 (-log<sub>10</sub>(p)<2) and log<sub>2</sub>(FC) > 0.18 or < - 0.18. Proteins with higher expression in wild-type (FC <0) in blue and with lower expression (FC>0) in red. For proteins with more than one differentially expressed feature, statistics refer to feature with largest FC effect is shown.

**Table S2:** Change in metabolites content in BECs exposed to *PbA*-IE detected in multiple targeting pathway analysis.

| Metabolite | p value | FC (exposed /unexposed) |
| --- | --- | --- |
| GLUTARATE | 7.48E-05 | 2.79 |
| PROPIOYLCARNITINE | 3.76E-03 | 2.28 |
| DEOXYRIBOSE | 4.38E-03 | 1.84 |
| OXOPROLINE | 2.47E-02 | 1.66 |
| ISOLEUCINE | 3.56E-02 | 1.58 |
| 4-ACETAMIDOBUTANOATE | 9.09E-03 | 1.53 |
| PANTOTHENATE | 3.55E-02 | 1.44 |
| CREATINE | 1.96E-02 | 1.39 |
| LACTATE | 4.53E-02 | 1.37 |
| ALANINE | 8.95E-03 | 1.36 |
| N,N,N-TRIMETHYLLYSINE | 2.94E-02 | 1.32 |
| HEXANOYLCARNITINE | 2.64E-02 | 1.31 |
| CIS-ACONITATE | 3.95E-02 | 1.31 |
| DIMETHYLARGININE (SDMA/ADMA) | 3.96E-02 | 1.29 |
| BUTYRYLCARNITINE | 2.25E-02 | 1.28 |
| BETAINE | 3.97E-02 | 1.19 |
| ASPARTATE | 2.43E-02 | 1.13 |
| THREONINE | 3.06E-02 | 0.81 |
| INOSINE MONOPHOSPHATE | 2.12E-02 | 0.69 |

P values of t-tests ( $P < 0.05$ ) and fold change (FC) comparing groups of three samples of exposed versus unexposed cells.

## FIGURE SUPPLEMENTAL LEGENDS

**Figure S1.**
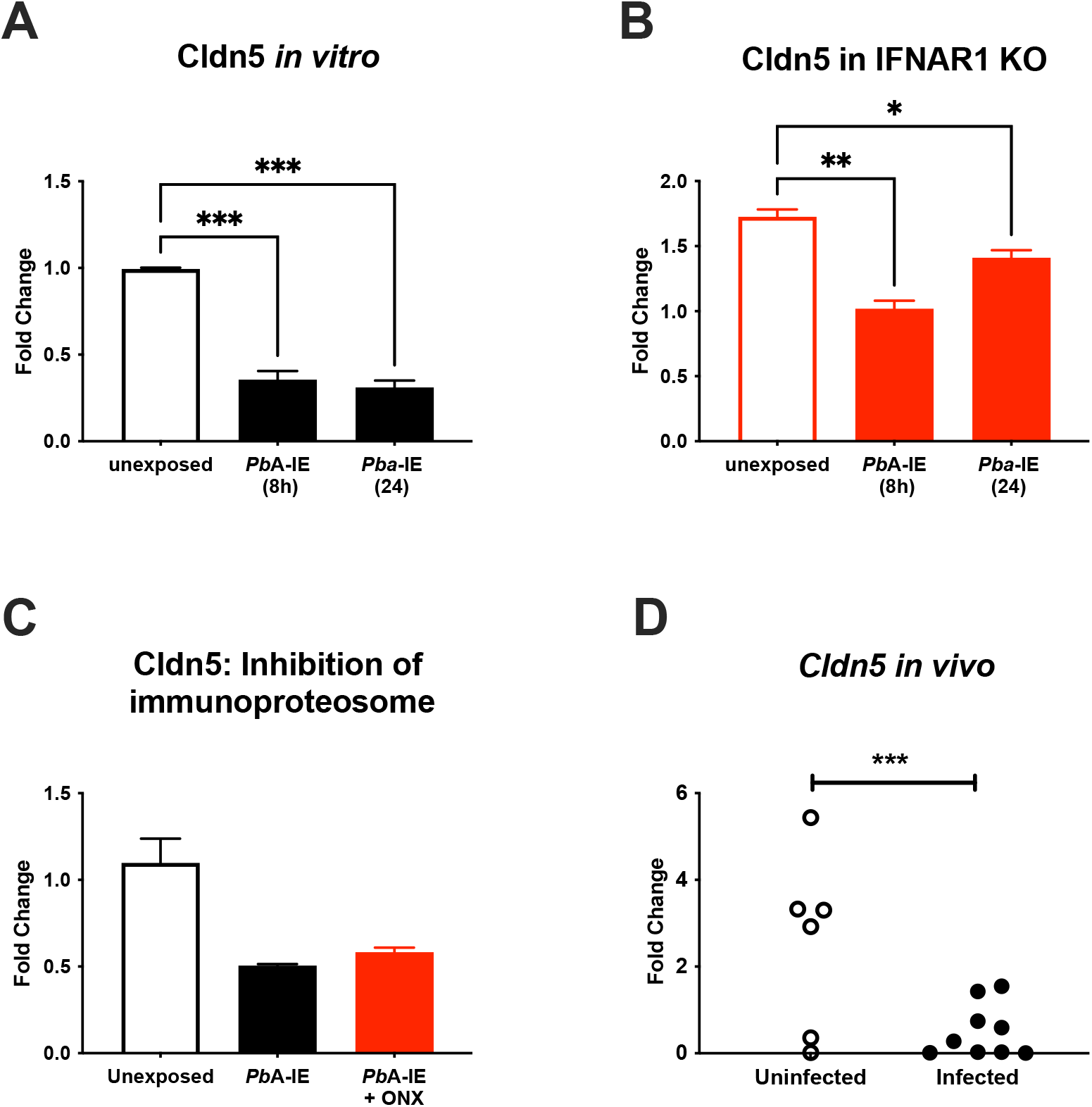
Claudin 5 gene expression in BECs is down-regulated upon exposure to *Pb*A-IE. Gene expression of *Cldn5* quantified in wild-type **(A)** and IFNAR1 KO BECs **(B)** at indicated time-points BECs 24 after exposure to *Pb*A-IE in the presence or absence of immunoproteasome inhibitor ONX-0914 (300 nM)(ONX)**(C)**. Gene expression of *Cldn5* quantified in brains of uninfected (n=6) and infected (n=9) mice **(D)**. Results of relative quantification gene expression are represented as fold change (2^ΔΔCT^) using unexposed BECs or uninfected mice as controls. Statistics: Significant results of pairwise comparisons in ANOVA tests are shown (*; p<0.05, **; p<0.01, ***; p <0.001).

**Figure S2.**
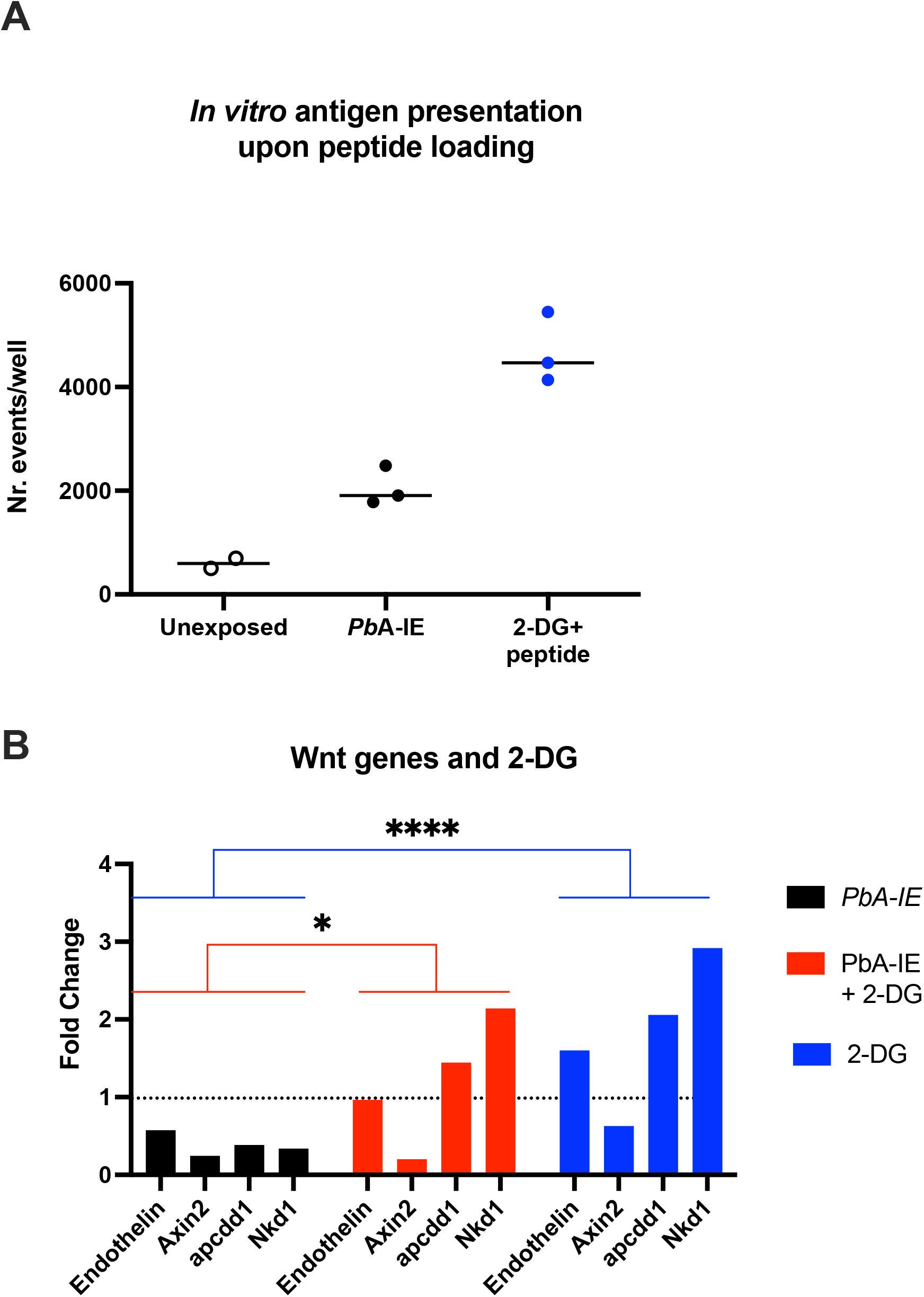
Effects of Glycolysis inhibition on antigen presentation of loaded peptides and expression of Wnt/*ß*-catenin target genes. Effects of inhibition of glycolysis by incubation of BEC cultures with 2-DG (10 mM) measured 24 h after exposure or not to *Pb*A-IE. **(A)** Antigen presentation of cognate peptide (SQLLNAKYL) (10 μM) loaded *in vitro* is not impaired by incubation with 2-DG. Results of pairwise comparisons in ANOVA tests (*; p<0.05, ***; p <0.001). **(B)** Gene expression of *ß*-catenin target genes *Edn1, Axin2, Nkdd1*, and *Apcdd1* as well as *cldn5* was quantified in wild-type BECs 24 h after exposure to *Pb*A-IE in the presence (red bars) or absence (black bars) of 2-DG or in presence of 2-DG (blue bars) alone. Relative quantification of gene expression is represented as fold change (2^ΔΔCT^) using unexposed BECs as controls (dashed line). Statistics: Results of pairwise group comparisons for 2-DG effect in 2-way ANOVA tests (*; p<0.05, ****; p <0.0001).

**Figure S3.**
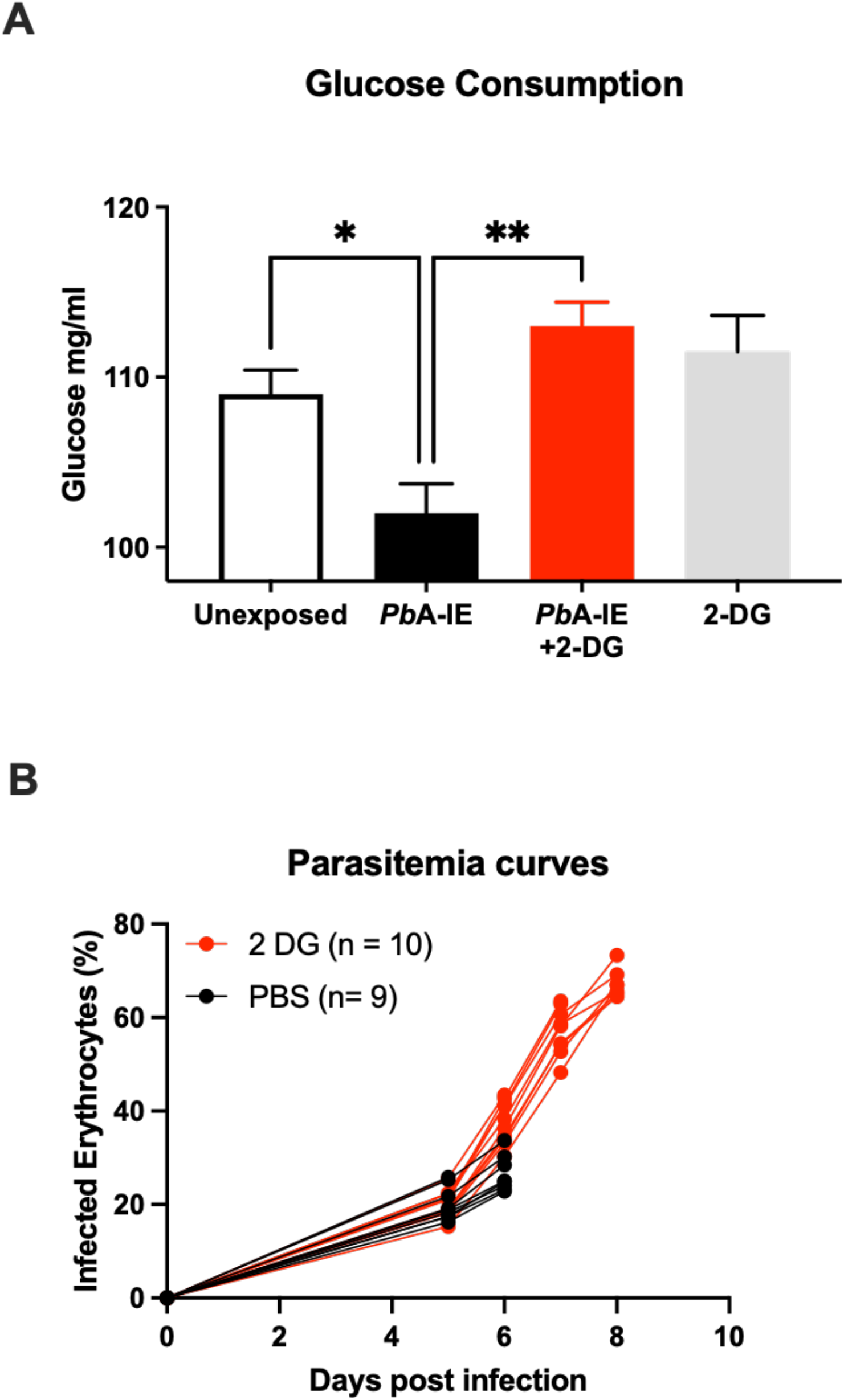
Glycolysis blockade: BECs glucose consumption *in vitro* and parasitemia *in vivo*. **(A)** Measurement of glucose in supernatants of BEC cultures exposed or not to *Pb*A-IE for 24 h in the presence or absence of 2-DG (10 mM). After 24-h exposure medium containing *Pb*A-IE was drained and cultures were replenished with fresh medium. Glucose measurements were performed 24 h after adding fresh medium. **(B)** Daily parasitemia measurements in individual mice infected with *Pb*A and treated (n=10) or not (n=9) with 2-DG injection (800 mg/kg) at day 4.

## Notes

### Competing Interest Statement

The authors have declared no competing interest.

### Summary of Updates

Abstract was improved and some references were corrected.

